# Leveraging current steering and the biophysics of spike generation for cellular-resolution electrical stimulation of neurons

**DOI:** 10.1101/2025.03.14.643392

**Authors:** Praful K. Vasireddy, Ramandeep S. Vilkhu, Amrith Lotlikar, Jeff B. Brown, A. J. Phillips, Alex R. Gogliettino, Madeline R. Hays, Claire Baum, Ethan J. Kato, Aviv Sharon, Pawel Hottowy, Alexander Sher, Alan M. Litke, Subhasish Mitra, Nishal P. Shah, E. J. Chichilnisky

## Abstract

Electrical stimulation at cellular resolution to restore the function of neural circuits is limited by the density of available electrode arrays. Although current steering with multi-electrode stimulation can be used to target cells between electrodes, it has not been proven for systematically targeting individual cells. We develop a framework for cellular-resolution current steering, leveraging the biophysics of electrically-evoked spike generation, and test its efficacy in isolated macaque and human retina. Currents were passed through three electrodes simultaneously using large-scale high-density microelectrode arrays, directly evoking single spikes in retinal ganglion cells. The currents combined either linearly or nonlinearly to drive spiking, depending on the geometry of the electrodes relative to the cell. These findings were captured by a biophysical model and by a simpler parametric model in which spikes can initiate at several sites on the cell membrane, and were leveraged to efficiently identify multi-electrode stimulation patterns that optimized cellular selectivity.

## Introduction

Electrical stimulation of neurons can be used to treat loss of nervous system function due to degeneration or trauma. Notable examples include deep brain stimulation for Parkinson’s disease [1] and cochlear implants for hearing loss [2]. However, a major unsolved challenge is the development of high-resolution electronic epiretinal implants for vision restoration. In retinal degenerative diseases such as age-related macular degeneration (200 million affected worldwide [3]) and retinitis pigmentosa (1.5 million affected worldwide [4]), electrical stimulation can bypass the damaged photoreceptor layer and directly evoke electrical activity in retinal ganglion cells (RGCs) which transmit visual information to the brain. Although the concept of the approach has been validated experimentally [5], epiretinal implants that target RGCs for electrical stimulation have not yet restored high acuity vision. One reason, revealed by decades of research on the neural code of the retina, is that natural vision is mediated by precise spatiotemporal spiking patterns of more than 20 RGC types that send distinct visual signals to various brain targets [6–8]. Because the distinct RGC types are intermixed on the surface of the retina, single-cell resolution is required to reproduce its neural code. However, this resolution is out of reach of existing and near-future epiretinal implants due to their coarse spatial resolution; consequently, present-day devices produce indiscriminate activation of RGCs of different types [5].

A potential way to more faithfully restore retinal signals is to use large-scale, high-density arrays of small electrodes, along with closed-loop targeting of distinct RGC types, to reproduce natural patterns of neural activity. Indeed, in *ex vivo* experiments using microelectrode array (MEA) stimulation and recording in peripheral macaque and human retinas [9,10], single-cell stimulation resolution is frequently achievable [11–14]. Unfortunately, this selectivity is not always achievable, and is more difficult in the central retina (which is responsible for high-resolution vision) because the spacing between RGCs is much smaller than typical MEA electrode pitch [12,15].

A possible way to overcome this limitation is *current steering*: simultaneous stimulation using multiple electrodes to more effectively target locations between electrodes. Current steering is well-established in neural stimulation at coarse spatial resolution [16–21], but only a few studies have examined the effect of multi-electrode stimulation at single-cell resolution [16,17,22] and our understanding of the biophysical underpinnings of current steering remains limited.

Moreover, previous work performed at cellular resolution required either exhaustive searches of all possible multi-electrode stimuli [17] or a specific geometric relationship between cells and electrodes to achieve selective activation [16,22], both of which are difficult to achieve in a clinical implant.

To overcome these challenges, we develop and experimentally validate a scalable, biophysically-inspired framework for understanding and predicting single-neuron responses to multi-electrode stimulation, and for efficiently searching the large space of all possible stimulation patterns to achieve selective stimulation to reproduce the neural code. Large-scale multi-electrode stimulation and recording from retinal ganglion cells using large-scale (512 electrodes) high-density (30 µm pitch) MEAs was performed in isolated macaque and human retinas, and the linear and non-linear current summation that produced short-latency directly-evoked spikes were interpreted using a biophysical model of neural activation. A simplified parametric activation model was then used for rapid identification of single-neuron responses to patterned electrical stimuli. To efficiently identify stimuli that could selectively activate a given cell while not activating others among the large number of possible multi-electrode stimulation patterns, data-driven regression models coupled with efficient sampling algorithms were used to speed calibration. These results leverage our biophysical understanding of electrical activation of neurons to enable current steering at cellular resolution and circuit scale, and are potentially relevant in other neural interface applications requiring cellular-resolution stimulation.

## Results

To probe the potential of current steering for precisely reproducing the neural code, multi-electrode stimulation and recording was deployed in an *ex vivo* preparation of human and macaque retinas using a 512-channel microelectrode array (MEA) system with 30 µm pitch. This approach was used to measure the activation of each targeted RGC (Fig. 1A) while also monitoring activity in several hundred nearby RGCs simultaneously. Triphasic, charge-balanced current pulses (50 μs phase duration, 0.1-1.8 μA second phase amplitude) were applied to three adjacent electrodes (an *electrode triplet*) simultaneously, and current amplitudes were varied independently on all three electrodes (Fig. 1B). This approach typically directly evoked a single spike in nearby RGCs with sub-millisecond latency [13,14]. Three-electrode stimulation was used because two-electrode stimulation can only steer electric fields in one dimension [16]. To characterize the stochastic activation of a targeted neuron (see [23]), each stimulus was repeated 20 times, evoked spikes were identified in the recorded traces using a custom spike sorting algorithm (see Methods), and an activation probability was calculated from these spikes (Fig. 1C). For each cell, the current pulse amplitude combinations in the three electrodes that produced 50% probability of an evoked spike were estimated and defined as the *thresholds* for activation, and the organization of these thresholds in the three-dimensional space of electrical stimuli was then examined (Fig. 1C) to test hypotheses about the nature of activation.

**Figure 1.**
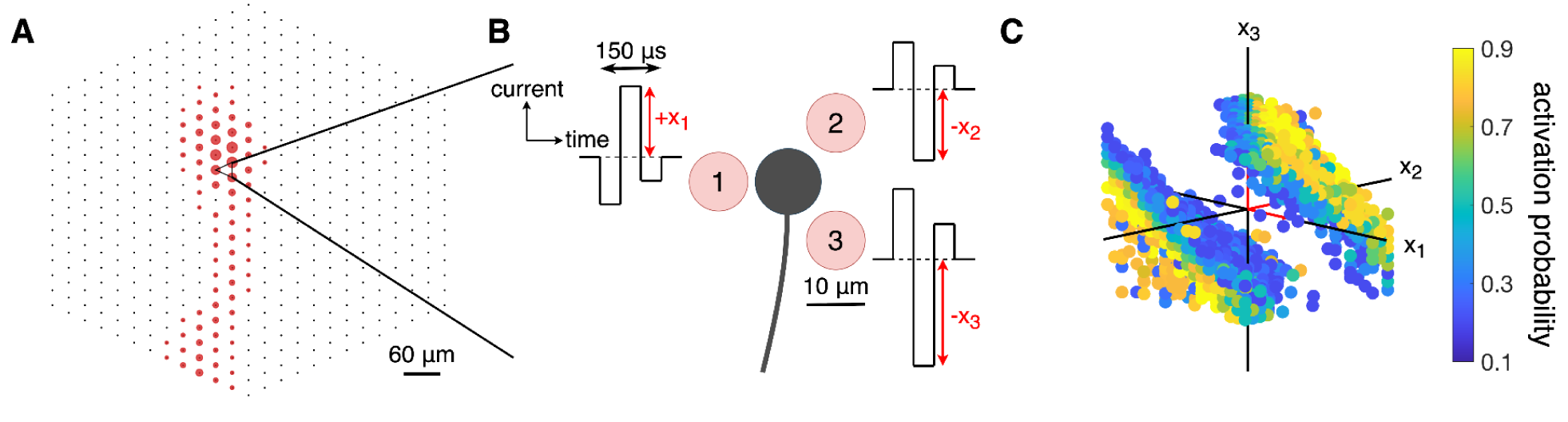
Illustration of the experimental data collection. ***A,*** Electrical image (see Methods) of a particular macaque peripheral OFF parasol cell recorded by the 512-channel MEA, with the three electrodes used for simultaneous electrical stimulation marked with a triangle. ***B,*** Zoom-in schematic view of the neuron (black) against the three stimulating electrodes in the MEA (light red) with electrode diameter and pitch of 10 µm and 30 µm, respectively. Triphasic, charge-balanced current pulses with 50 μs phase duration and relative phase amplitudes of (2:-3:1) are applied to all three electrodes simultaneously. The second phase amplitude (a scalar) is used for visualization in subsequent plots and is varied independently across the three electrodes. ***C,*** Visualization of single-neuron responses to combinations of stimulation currents on three electrodes. Activation probabilities are calculated by averaging spiking responses across repeated trials of the same stimulus current combination. Red sections of axes mark the positive direction and are of length 1 μA. Only intermediate probabilities with 0<p<1 are shown for visual clarity.

### Neurons exhibit linear and nonlinear responses to multi-electrode stimulation

In the simplest case, neural activation could be evoked by a linear summation of currents from the three electrodes. Specifically, if the currents (*x*_1_, *x*_2_, *x*_3_) from the three electrodes sum linearly to evoke a spike, then the collection of stimuli producing an activation probability of 50% – activation *thresholds* – must satisfy the relation σ(*w*_0_+*w*_1_*x*_1_+*w*_2_*x*_2_+*w*_3_*x*_3_)=0.5, where σ() represents the potentially nonlinear relationship between current amplitude and spike probability. In the present work, σ() is a sigmoid, consistent with approximately Gaussian membrane voltage noise and consequent stochastic spike production [13,23]. From the above equation, linear summation predicts that the collection of measured thresholds (*x*_1_, *x*_2_, *x*_3_) form a plane in the space of three-electrode stimuli.

The results from 22 tests of summation on 65 neurons, performed using RGCs of several types recorded at several retinal eccentricities in human and macaque retinas, revealed both linear and nonlinear responses to electrical stimulation. In some cases, the thresholds formed approximately planar surfaces (Fig. 2A-D). Because a charge-balanced stimulation was used, anodic-peak stimuli (positive current values in Fig. 2) often elicited spiking at similar threshold levels as cathodic-peak stimuli (negative current values in Fig. 2), resulting in a second planar threshold surface (note that the two planes were not necessarily symmetric about the origin, in part because of the asymmetry between cathodic and anodic phases of the stimulating pulse; see Fig. 2A,D, Methods). This two-plane geometry thus indicates that, in some cases, current from the three electrodes summed approximately linearly to evoke spikes. In other cases, the collection of activation thresholds formed a curved surface (Fig. 2E-H), revealing nonlinear summation of currents across electrodes. The nonlinearity was sometimes sufficiently strong that the surface was closed, even within the limited range of current levels tested (Fig. 2E,G,H). This structure confirms and extends previous findings with two-electrode stimulation (see [17]).

**Figure 2.**
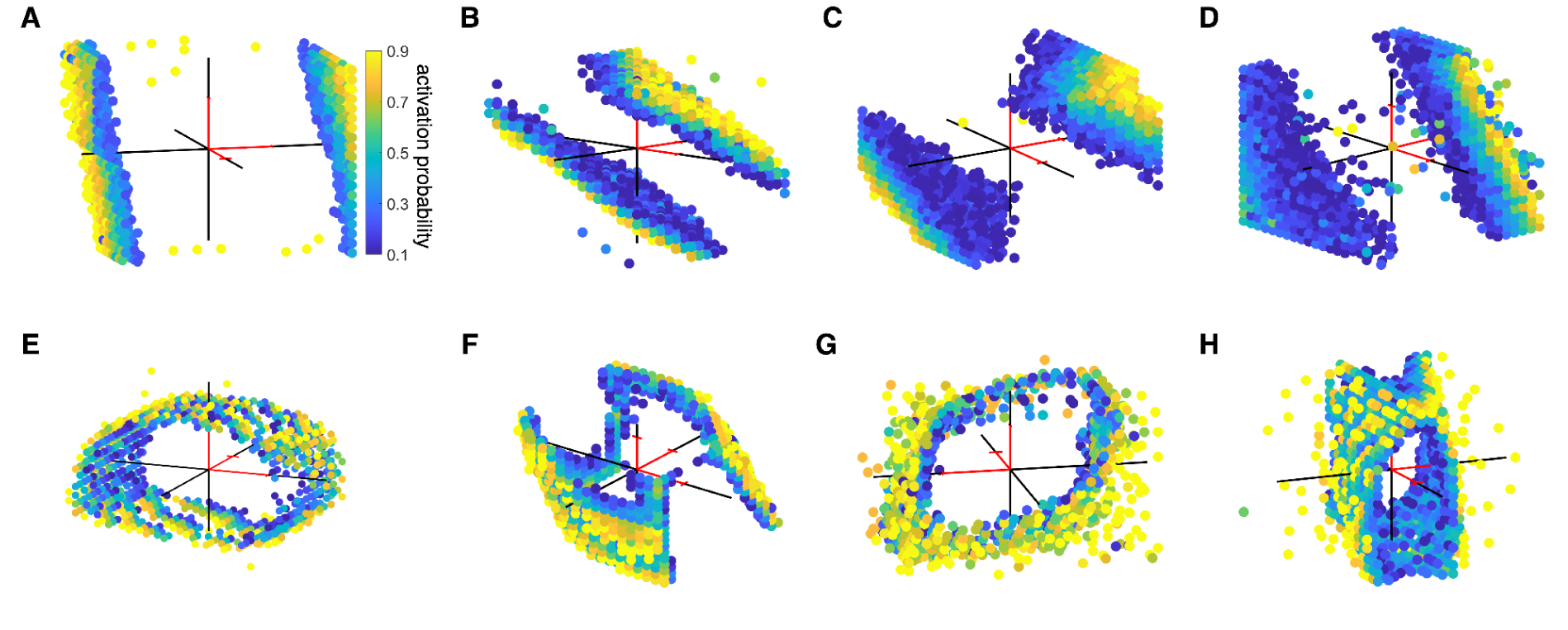
Experimental data for three-electrode stimulation of individual neurons. Macaque and human retinal ganglion cells of different cell types from various retinal preparations are stimulated with three electrodes simultaneously at independently varying current amplitudes (*x*_1_, *x*_2_, *x*_3_), and their response profiles are measured. The stimulating current on each electrode varied in second phase amplitude from −1.8 μA to 1.8 μA. Each panel represents the aggregated response probabilities collected from performing 160,000 independent stimulations (30 minutes of stimulation). The full dataset comprised 65 neurons, each of which displayed a threshold surface, and the panels included here are representative of the variety of surfaces observed. Red sections of axes mark the positive direction and are of length 1 μA. Only intermediate activation probabilities with 0<p<1 are shown for visual clarity of the threshold surface. ***A,*** Macaque central ON parasol. ***B,*** Macaque peripheral OFF parasol. ***C,*** Human central ON parasol. ***D,*** Human peripheral OFF midget. ***E,*** Macaque peripheral ON parasol. ***F,*** Macaque peripheral ON parasol. ***G,*** Macaque peripheral ON parasol. ***H,*** Macaque central OFF parasol.

### Biophysical simulations explain linear and nonlinear responses to multi-electrode stimuli

To test whether the linear and nonlinear activation surfaces observed with different electrode geometries were consistent with known properties of RGCs, a biophysical simulation was developed with cellular properties and MEA geometry approximately matching the experiments (see Methods; Fig. 3A). The biophysical model was developed in the NEURON simulation environment [24] and consisted of dendritic, somatic, and axonal compartments. The ion channel parameters in each compartment and cytoplasmic (axial) resistivity were set to match those of primate RGCs, and the temperature was chosen to match experimental conditions. The stimulating electrode geometry and pulse waveforms were also arranged to match the experiments (see [25] for details).

**Figure 3.**
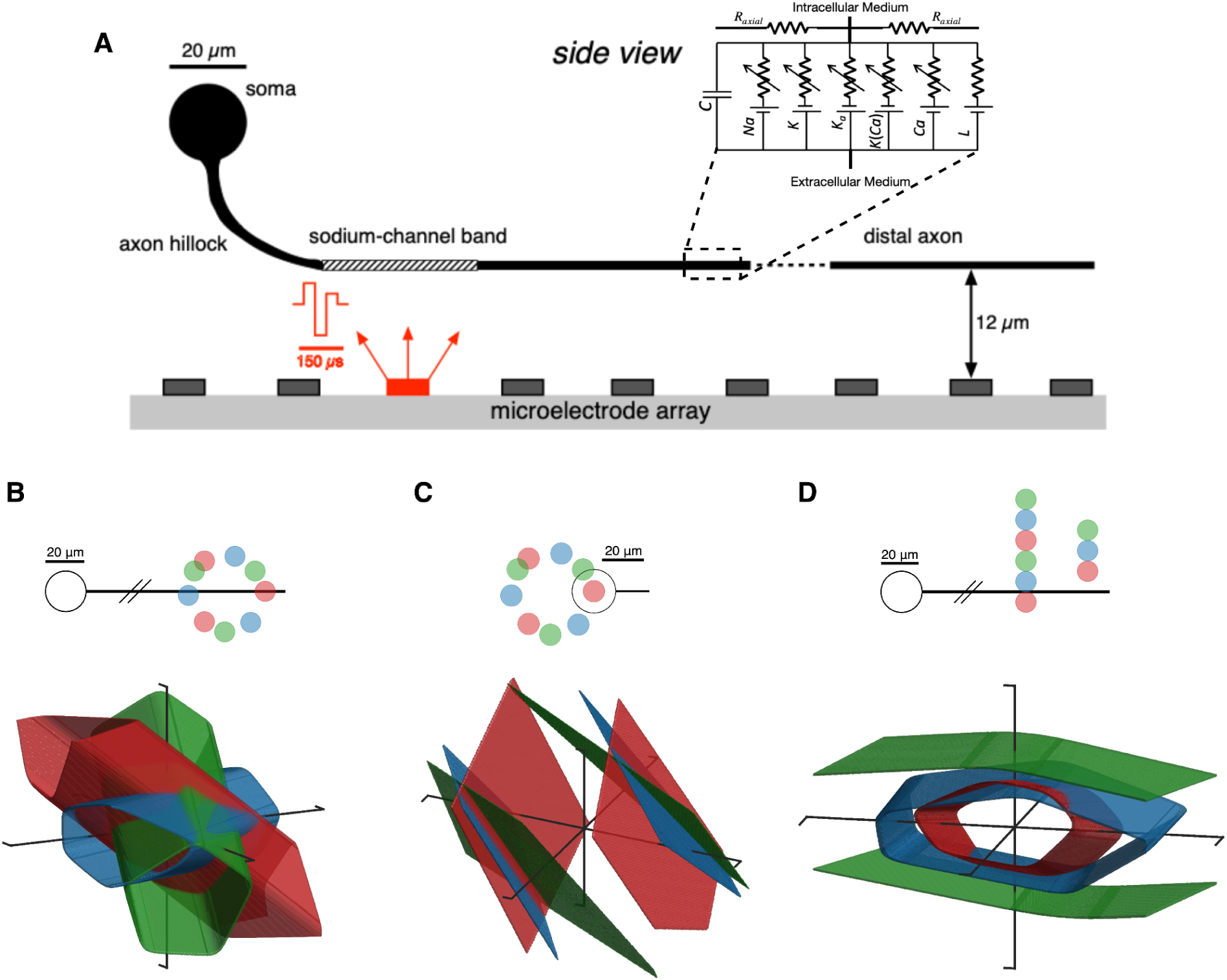
Biophysical simulation responses for three-electrode stimulation. ***A,*** Side view of the model cell showing the various modeled compartments and the bend in the RGC axon as it extends from the soma to the nerve fiber layer, where the planar MEA is positioned, matching experimental conditions. Zoom-in shows the equivalent circuit for each segment of the biophysical model, with conductances labeled for sodium (Na), potassium (K), a-type potassium (K_a_), calcium-gated potassium (K(Ca)), calcium (Ca), and passive leak (L) channels. Charge-balanced triphasic stimulus with relative phase amplitudes of (2:-3:1) supplied through a single electrode (red). Dendrites not shown. ***B-D,*** Colored surfaces represent the 50% activation probability boundary. Schematic headings show a top-down (x-y) view of the relative geometry between the neuron and stimulating electrodes, with the color of the electrodes corresponding to the color of the 50% activation probability boundary from stimulating with those electrodes. The electrodes lie on a fixed-z plane 12 μm above the distal axon and 40 μm above the soma center. ***B,*** Stimulation at the distal axon with varying rotation angle. ***C,*** Stimulation near the somatic compartment with varying rotation angle. ***D,*** Stimulation at the distal axon with varying transverse offset.

The biophysical model cell exhibited linear or nonlinear summation, with the degree of the nonlinearity determined by the electrode-cell geometry that governed the site(s) in the cell where the spike was initiated. Specifically, stimulating electrodes placed near the cell body produced nearly linear responses (planar surfaces) (Fig. 3C) while electrodes placed near the distal axon produced nonlinear responses (curved surfaces) (Fig. 3B,D). Stimulating electrodes placed near the soma most often initiated spikes at the sodium-channel band, a location on the proximal axon with the highest density of voltage-gated sodium channels, whereas electrodes placed near the distal axon could initiate spikes at several locations along the axon [25]. The rotation angle of the electrodes relative to the neuron affected the orientation of the threshold surfaces but did not significantly alter the degree of nonlinearity (Fig. 3B,C). Increasing the distance between the electrodes and the cell produced increasingly linear response surfaces (Fig. 3D), because electrodes had electrical access to fewer distinct spike initiation sites within a given stimulating current amplitude range [25]. Similar findings were observed using biphasic rather than triphasic stimulating current pulses (data not shown). These features are qualitatively consistent with multi-site activation of RGCs demonstrated in previous work [25].

### A simple parametric activation model captures linear and nonlinear summation

To test whether linear and nonlinear activation can be understood and harnessed to target neurons for electrical stimulation, a parametric model of activation was developed. This model, which we refer to as the *linear-OR model*, is biophysically motivated, can be rapidly fitted to experimental data, and can be used to accurately predict single-neuron responses to untested multi-electrode stimuli.

The linear-OR model is designed to approximate the probability of a spike that can be generated at any of several distinct sites in a cell [17,25]. In the model, each site linearly sums the applied stimulation current on many electrodes. This sum in turn controls the probability of producing a spike at that site via a sigmoidal relationship, which reflects membrane voltage noise. The probability of a spike in the cell is the probability that any site produced a spike. The linear-OR model is mathematically described by:

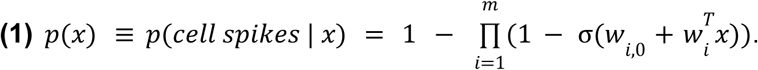

Here, *x* is the vector of stimulation currents applied to a collection of electrodes, *m* is the number of distinct activation sites, *σ*() is the sigmoid function that controls activation probability [13], and *w_i,0_* and *w_i_* are biases and weights for each site. Equation 1 represents the complement of the probability that none of the *m* initiation sites are activated, or equivalently, the probability that *any* of the initiation sites is activated. Simply put, this model represents a logical OR operation between *m* linear spike initiation sites. For *m* = 1, fitting the linear-OR model reduces to fitting a logistic regression model, while increasing values of *m* allow fits to increasingly nonlinear data.

To test the linear-OR model on experimental data, the three-electrode activation threshold surfaces were fitted with the linear weights and biases as free parameters (see Methods). The fitted model accurately captured both linear (Fig. 4B) and nonlinear (Fig. 4A,C) activation surfaces. The mean absolute error between model predictions and observed activation probability was significantly lower than that of a linear model (see Methods; Fig. 4D-F).

**Figure 4.**
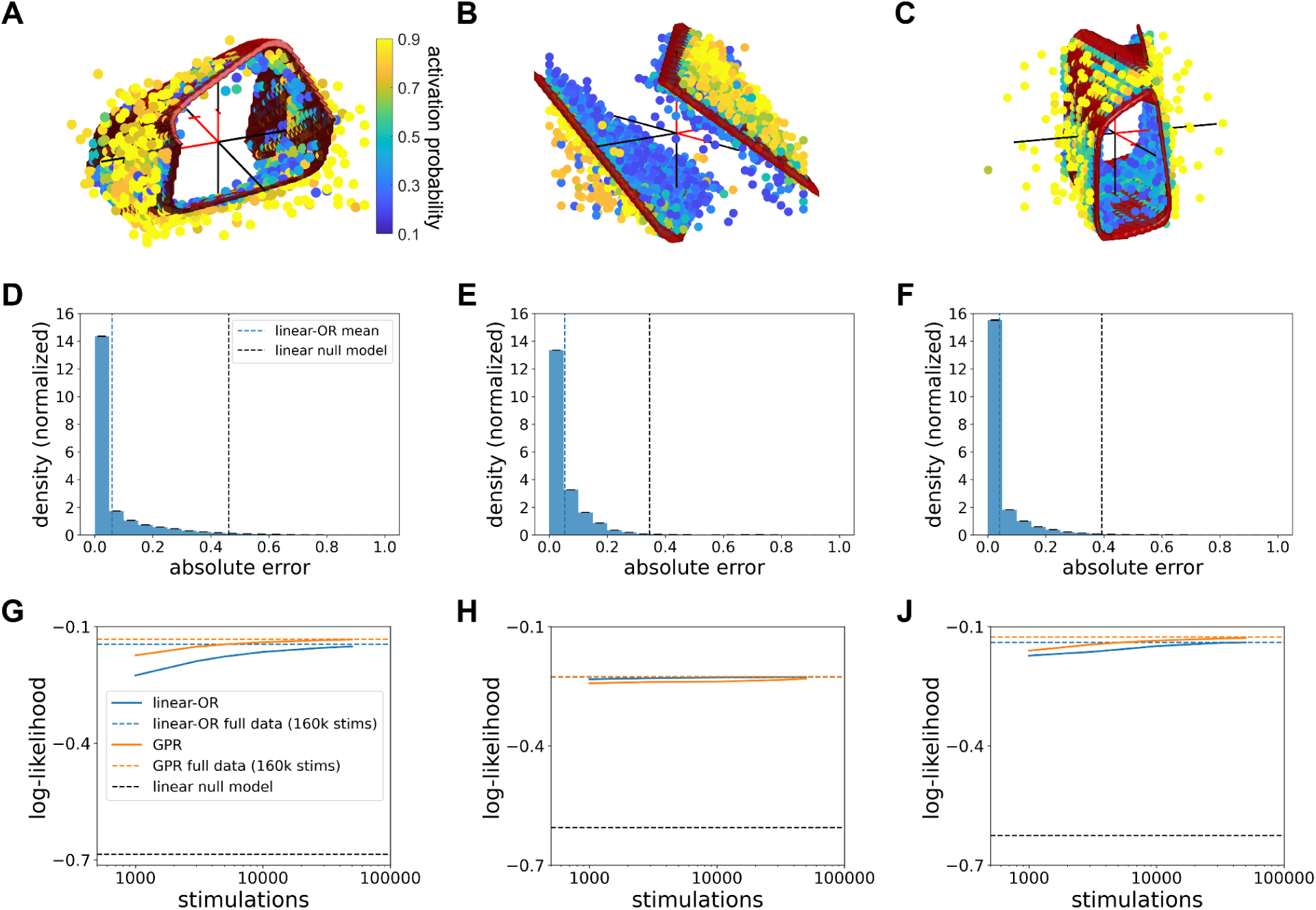
Linear-OR activation model fitting to experimental data. Red surfaces represent the 50% activation probability surface predicted by the linear-OR model fit to the full experimental data. Red sections of axes mark the positive direction and are of length 1 μA. ***A,*** Nonlinear three-electrode stimulation experimental data (macaque peripheral ON parasol) with linear-OR model 50% surface for *m* = 4 activation sites. ***B,*** Linear three-electrode stimulation experimental data (macaque peripheral OFF parasol) with linear-OR model 50% surface for *m* = 2 activation sites. ***C,*** Nonlinear three-electrode stimulation experimental data (macaque central OFF parasol) with linear-OR model 50% surface for *m* = 4 activation sites. ***D–F,*** Distribution of error between fitted linear-OR model predictions and experimental data, for neurons from *A–C*. Blue vertical dashed line represents the mean absolute error achieved by the linear-OR model, while black vertical dashed line is the mean absolute error achieved using a linear null model. 100 refittings were used for error bars. ***G–I,*** Log-likelihood between predictions from linear-OR model or Gaussian process regression (GPR) model and experimental data as a function of number of calibrating stimulations, for neurons from *A–C*. Colored horizontal dashed lines represent the log-likelihood level achieved by fitting to the full data (160,000 stimulations), and black horizontal dashed line represents the log-likelihood level achieved using a linear null model. 100 samples were used for confidence intervals.

The linear-OR model was also tested by fitting to threshold surfaces produced using biophysical simulations with realistic neuron-cell geometries. Simulated surfaces with electrode geometries producing a range of linear (Fig. 5B) to highly nonlinear (Fig. 5A,C) behaviors were accurately fitted by the model. As a test of generalization to larger numbers of electrodes, simulations of four-electrode (Fig. 5E) and five-electrode stimulation (Fig. 5F) were performed. Three-dimensional sections of activation surfaces again revealed accurate model performance (see Extended Data Fig. 1).

**Figure 5.**
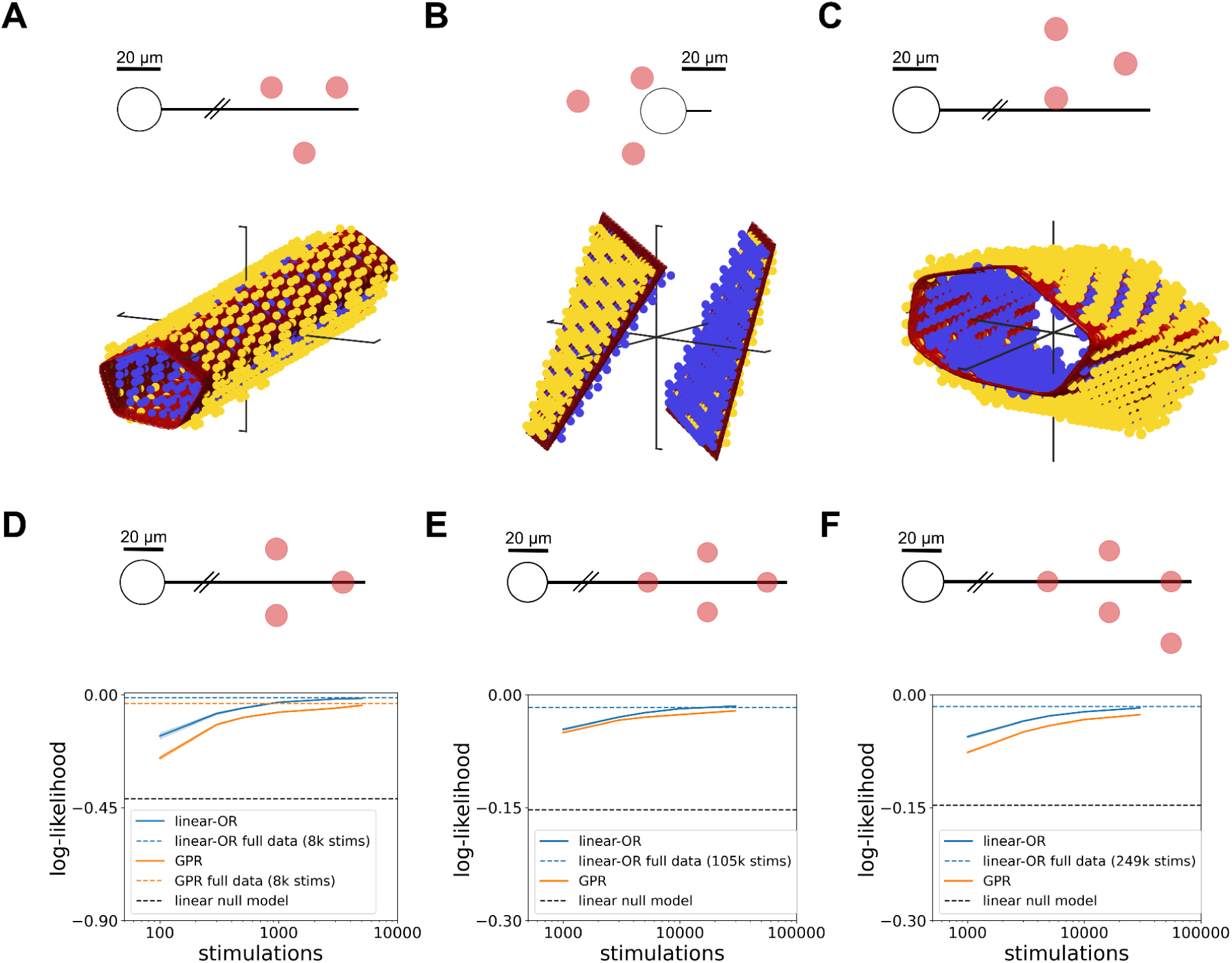
Linear-OR activation model fitting to biophysical simulation data. Red surfaces represent the 50% activation probability surface predicted by the linear-OR model fit to the biophysical simulation data. Schematic headings show a top-down (x-y) view of the electrodes (red) relative to the neuron (black). The electrodes lie on a fixed-z plane 12 μm above the distal axon and 40 μm above the soma center. ***A,*** Nonlinear three-electrode stimulation biophysical simulation data with linear-OR model 50% surface for *m* = 8 activation sites. ***B,*** Linear three-electrode stimulation biophysical simulation data with linear-OR model 50% surface for *m* = 2 activation sites. ***C,*** Nonlinear three-electrode stimulation biophysical simulation data with linear-OR model 50% surface for *m* = 8 activation sites. ***D-F,*** Log-likelihood between predictions from linear-OR model or Gaussian process regression (GPR) model and biophysical simulation data as a function of number of calibrating stimulations, for three-, four-, and five-electrode stimulation. GPR full data fits not included for ***E,F*** due to computational limitations. 100 samples were used for confidence intervals.

To provide a comparison to a more flexible and general (but less interpretable) model, Gaussian process regression (GPR) was tested for both experimental and simulated data [26,27] (see Methods). In linear experimental data (Fig. 4B,H), the linear-OR model slightly outperformed the GPR model; in nonlinear data, the two models performed similarly (Fig. 4A,C,G,J), although the linear-OR model typically required more data to achieve maximum performance (Fig. 4G,J). In simulated data with no biological or measurement noise, the linear-OR model consistently outperformed GPR even when the number of stimulating electrodes was increased (Fig. 5D-F).

### Multi-electrode stimulation enables significant improvements in selectivity

A central goal of current steering is to selectively activate targeted neurons without activating others, to most effectively match the neural code. In the case of triplet stimulation, achieving selectivity corresponds to finding regions of the three-dimensional space of current amplitudes that are outside the threshold surface of a target cell but within the threshold surface of non-target cells (Fig. 6A).

**Figure 6.**
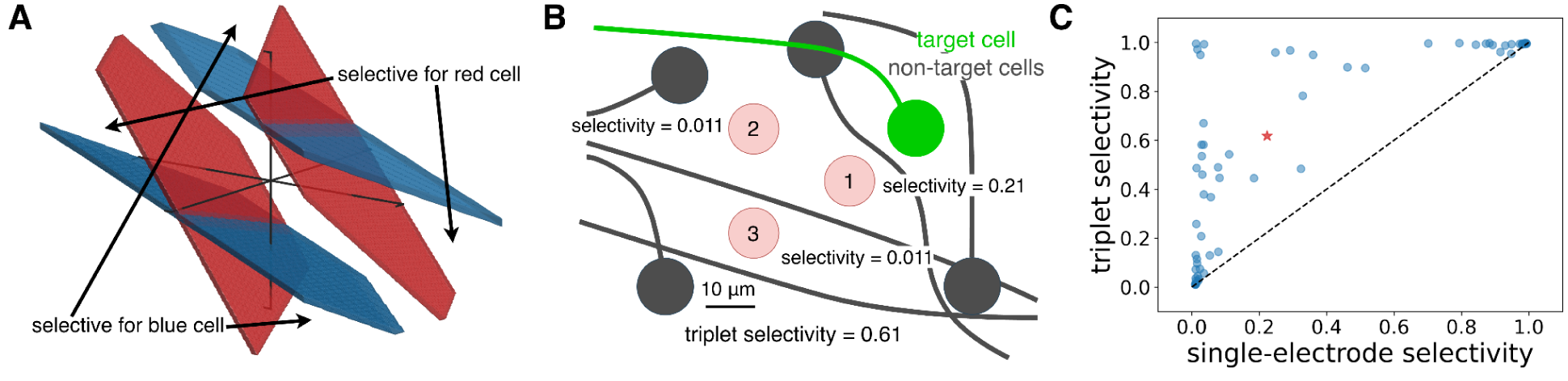
Selectivity obtained from multi-electrode stimulation. ***A,*** Schematic of 50% activation probability surfaces (red and blue surfaces) for two neurons stimulated by the same electrode triplet, with selective regions in stimulus space for each cell indicated. Surfaces were extracted from measurements from macaque peripheral RGCs. ***B,*** Schematic of selectivity improvement for an example with seven macaque central RGCs. Red numbered circles denote electrodes, with relative positions of non-target cells (gray) and target cell (green) shown. Each electrode is labelled with its maximum achievable selectivity index using single-electrode stimulation. ***C,*** Maximum selectivity index obtained with triplet stimulation compared to maximum selectivity index obtained with single-electrode stimulation on the same set of electrodes, accumulated across 65 RGCs from human and macaque retinal preparations. Red star indicates the example shown in panel B.

To probe the effectiveness of triplet stimulation, a metric of selectivity was developed. For a collection of neurons *c*_1_,…,*c*_K_ that can be activated using electrical stimulus *x*, the *selectivity index*, *S_i,x_*, for a given target cell, *c_i_* , with this stimulus is given by:

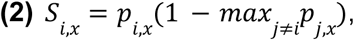

where *p_i,x_* is the activation probability of the target cell *c_i_* using stimulus *x,* and *p_j,x_*is the activation probability of a non-target cell *c_j_* using stimulus *x*. The non-target cell *c_j_* is selected to have the maximum activation of all non-target cells, i.e. the worst case. The metric ranges from 0 to 1, with 1 indicating perfect selectivity. For example, a stimulus that elicits 70% probability of spiking in the target cell and 30% probability of spiking in the worst-case non-target cell yields a selectivity index of 0.49. The maximum of the selectivity index across all stimuli using a specific electrode triplet was measured, including the maximum selectivity obtained with each electrode individually. By definition triplet stimulation is always at least as selective as single-electrode stimulation because single-electrode stimulation is a special case of triplet stimulation.

Triplet stimulation frequently provided a significant improvement in selectivity over single-electrode stimulation. For example, in the case of a macaque central OFF parasol RGC (Fig. 6B), selectivity indices 0.21, 0.011, and 0.011 were obtained using each of the three electrodes individually, but the optimal three-electrode stimulation pattern yielded selectivity of 0.61. To assess the visual significance of this enhanced selectivity, a data-driven simulation of visual reconstruction from evoked spikes was performed using the framework from [12,28] for an experimentally recorded population of RGCs [28]. The reconstruction error was significantly reduced by increasing the single-neuron selectivity across the population from an average of 0.2 to an average of 0.6, and the error was most quickly reduced as selectivity was increased from 0.6 to 1.0 (not shown).

The results accumulated across 65 human and macaque RGCs tested with three-electrode vs. single-electrode stimulation suggest that current steering can be very effective for increasing selectivity (Fig. 6C). Using only single-electrode stimulation, 22/65 cells (33.8%) had a selectivity index greater than 0.5, compared to 37/65 cells (56.9%) using triplet stimulation. The remaining 28/65 cells (43.1%) tested could not be selectively activated with either single- or three-electrode stimulation using the 22 electrode triplets that were experimentally tested.

However, this does not necessarily mean the selectivity of these remaining neurons cannot be improved with multi-electrode stimulation: many triplets of electrodes were not tested due to experimental time constraints. This limitation highlights the need for efficient sampling procedures to rapidly identify selective stimulation patterns on many groups of three electrodes (see below).

### Rapid identification of selective multi-electrode stimuli using data-driven models

The effectiveness of current steering for selectivity relies on efficient calibration, because it is unknown *a priori* which groups of electrodes will selectively stimulate a neuron, and because exhaustive calibration with a large-scale MEA is likely impractical in a clinical setting. Therefore, a method was developed to rapidly identify selective stimuli, and its performance was evaluated.

First, a measure of aggregate selectivity was defined as the selectivity index obtained considering the most selectively activatable cell as the target (see Methods Eq. 5). Then, the aggregate selectivity across all tested three-electrode stimuli was computed (see Fig. 7A), defining the space that must be searched to determine the most selective stimuli.

**Figure 7.**
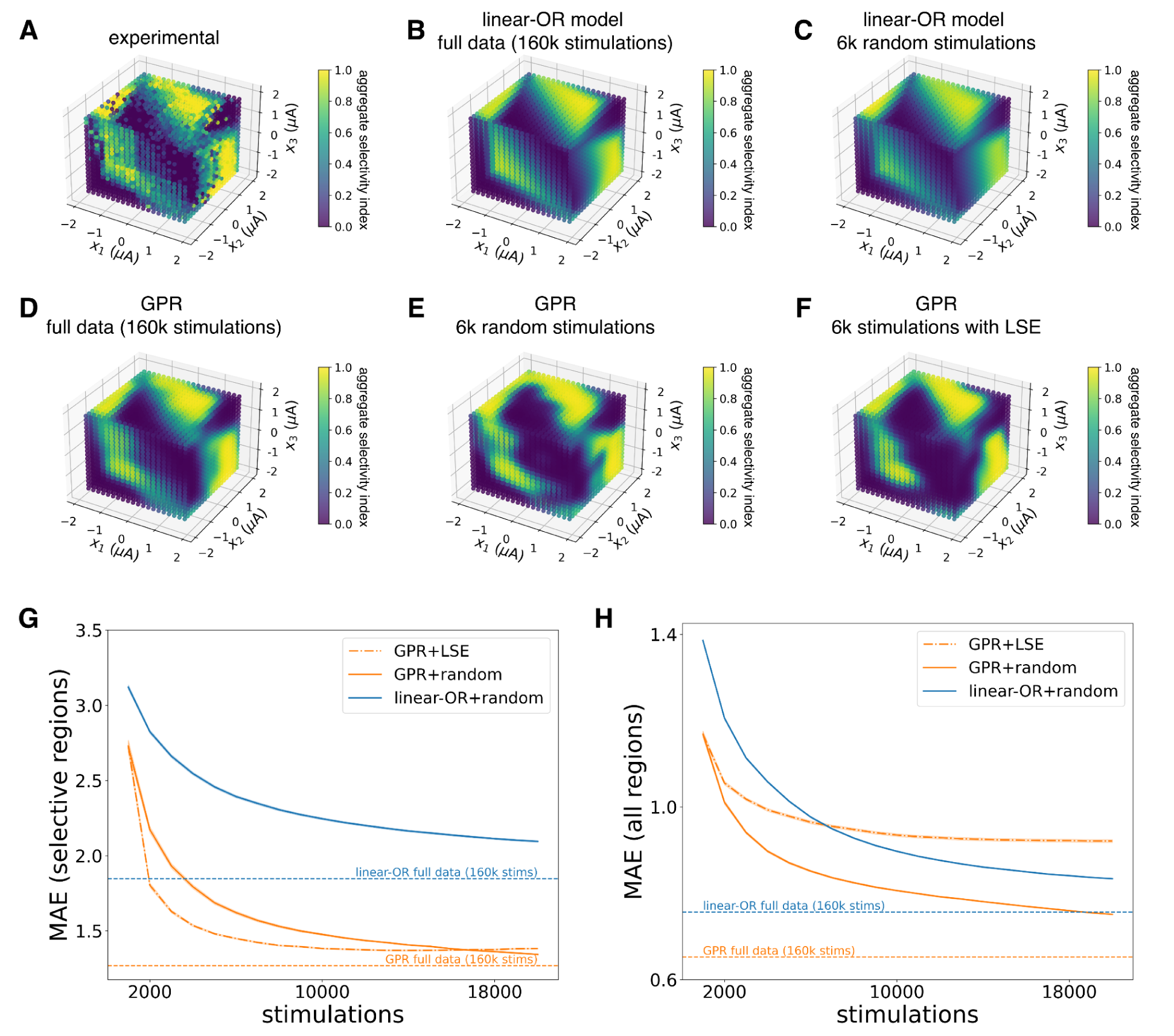
Aggregate selectivity landscapes fitted with linear-OR model and Gaussian process regression (GPR). ***A,*** Experimental aggregate selectivity profile for three-electrode stimulation, aggregated across three macaque central RGCs stimulated by the same electrode triplet. Remaining panels show fitted selectivity predictions using ***B,*** linear-OR model, exhaustive data (160k calibrating stimulations), ***C,*** linear-OR model, 6k randomly sampled calibrating stimulations, ***D,*** GPR, exhaustive data (160k calibrating stimulations), ***E,*** GPR, 6k randomly sampled calibrating stimulations, ***F,*** GPR, 6k stimulations sampled with the GPR+LSE procedure. ***G,*** Mean absolute error (MAE) in log-odds units between model predictions and experimental data in high selectivity regions (the union of regions with selectivity index greater than 0.9 as estimated by the model and as measured by experimental data) as a function of number of calibrating stimulations, with GPR+LSE sampling compared to GPR with uniform random sampling. MAE of the linear-OR model fitted to randomly subsampled data is also shown. 100 restarts were used for confidence intervals. ***H,*** Mean absolute error (MAE) in log-odds units between model predictions and experimental data in all regions as a function of number of calibrating stimulations, with GPR+LSE sampling compared to GPR with uniform random sampling. MAE of the linear-OR model fitted to randomly subsampled data is also shown. 100 restarts were used for confidence intervals.

An obvious candidate approach to rapidly identify selective stimuli would be to use the linear-OR activation model fitted to limited experimental data and evaluated on all possible stimuli in this space. To test this possibility, the linear-OR model was fitted to either the entire data set or a 3.75% subset of the data. The model captured the large-scale features of the measured selectivity landscape when it was fitted with all the data (Fig. 7B,H), and its predictions changed little even with significant subsampling (Fig. 7C). However, some of the details of the complex, experimentally determined selectivity landscape (Fig. 7A) were not fully captured by the linear-OR model, especially in the most relevant highly selective regions (see Fig. 7G).

As a more flexible alternative, a model of selectivity based on GPR was tested by again fitting the selectivity landscape with a subset of measured responses and predicting selective stimuli. Compared to the linear-OR model, GPR was able to capture more precise details of the selectivity landscape using all the recorded data (Fig. 7D,G). However, due to the flexibility of the GPR model, random subsampling often led to less accurate fits in highly selective regions (Fig. 7E,G). This finding suggests that GPR fits could be improved by adaptive sampling procedures that preferentially explore informative regions of the selectivity landscape.

Accordingly, GPR was used to develop an efficient closed-loop sampling scheme based on a method known as level set estimation (LSE; see Methods). Briefly, performing GPR provides a prediction of the activation probability along with a metric of uncertainty at every combination of stimulation currents (see Methods). The LSE scheme leverages the uncertainty metric and prioritizes sampling combinations of currents that are both uncertain and could yield high selectivity, and deprioritizes sampling combinations of currents with low uncertainty or known extreme selectivity (high or low). Using this sampling scheme (denoted as GPR+LSE), highly selective regions were well estimated even with significant subsampling (Fig. 7F,G).

The GPR+LSE framework substantially reduced the total time required to accurately determine selective stimulation patterns. Highly selective regions (selectivity index > 0.9) were estimated more accurately and with fewer measurements using the GPR+LSE procedure than with random sampling (Fig. 7G, Extended Data Fig. 2). In a specific example with three macaque RGCs in the central retina stimulated by a specific group of three electrodes, sampling with GPR+LSE required ∼10,000 calibrating stimulations to achieve accuracy within 10% of the accuracy obtained by performing GPR on the exhaustive data, compared to ∼15,000 stimulations required to reach the same accuracy level with random sampling (Fig. 7G). Both procedures showed substantial improvement over using the linear-OR model, requiring ∼2,000 stimulations with GPR+LSE and ∼3,000 stimulations with random sampling to achieve the same accuracy as the linear-OR model fitted to the exhaustive data (160k stimulations; Fig. 7G).

However, because the GPR+LSE procedure prioritized estimating selective regions, unselective regions were more poorly estimated than with random sampling, leading to higher error across the whole selectivity landscape (Fig. 7H). This drawback was alleviated by a hybrid scheme that mixed LSE sampling with uniform random sampling (not shown). In sum, several approaches can be used to efficiently characterize triplet selectivity, and the most efficient procedure depends on the goal of the measurement.

## Discussion

Experimental investigation of cellular-resolution current steering with groups of three electrodes yielded two major findings. First, a simple biophysically interpretable model of multi-site activation captured the main trends in linear and nonlinear neural activation, permitting a simple and accurate prediction of neural responses to arbitrary electrical stimulus patterns with three electrodes. Second, the ability to selectively activate one neuron without activating others was significantly enhanced by current steering, and empirical models permitted efficient sampling to identify the most selective stimuli. These findings suggest that current steering could be used to more precisely reproduce the neural code at cellular resolution in future electronic implants.

The linear-OR model of neural activation with multiple electrodes provided a biophysically motivated way to understand the linear and nonlinear properties of electrical activation with relatively few assumptions and measurements. Biological tissue conductivities are approximately linear at the frequencies used here [29,30], meaning that a scalar linear weight can be assigned between the stimulating current on an electrode and the resulting change in extracellular voltage at a specific region on a passive cell membrane. Other functions of the membrane potential relevant to eliciting spikes extracellularly, such as its second directional derivative along the cell membrane (activating function; [31]), are also linear in the stimulating current. However, unlike normal physiological spike initiation, which occurs at the sodium channel band in the proximal axon [32], extracellular stimulation of neurons may initiate spikes at various locations on the cell membrane depending on the geometric arrangement of the stimulating electrodes relative to the cell. This fact is seen in the inadvertent stimulation of RGC axons with epiretinal implants [11]. The ability to generate a spike at each of several locations leads to nonlinear neural responses to multi-electrode stimuli, even if spike initiation at a single location is linear [25]. Importantly, nonlinear interactions are clearly visible using low currents and short distances between electrodes and neurons, but are much more difficult to observe with distant electrodes that effectively target the same site [25,33].

Although the parametric linear-OR model accurately predicted and fitted experimental data, the numerical values of the fitted weights were only easily physiologically interpretable in linear cases with *m* ≤ 2, for which the magnitude of the weight was inversely related to the distance from the stimulating electrode to the activation site (data not shown). In contrast, the physiological interpretation of the linear-OR model parameters is more difficult in nonlinear cases with *m* > 3. Nonlinearities arise from stimulating the uniformly excitable axon fiber, and the linear-OR model approximates this underlying continuous excitability with a large number of discrete activation sites (typically 2 < *m* ≤ 10 for three-electrode stimulation), limiting physical interpretability. However, the response surface contains some geometric information about the neuron, such as the direction of the axon relative to the rotation of the stimulating electrodes (see Fig. 3B) or the offset of the electrodes relative to the axon (see Fig. 3D). Accordingly, it may be possible to learn this information from the fitted linear-OR model parameters alone, although analytical solutions may not exist.

The linear-OR model fitted complex nonlinear data accurately, and accurately captured data obtained using four or five stimulating electrodes. With more stimulating electrodes, the complexity of the evoked response and the number of possible stimulation patterns grows. The biophysical properties that underpin the linear-OR model and its low computational complexity for fitting (*O*(*N*) with *N* the number of data points, compared to other nonlinear fitting methods such as GPR which are *O*(*N*^3^)), allow it to effectively scale to more stimulating electrodes.

However, increasing the number of stimulating electrodes may not improve selectivity, because each added stimulating electrode typically causes activation of additional cells. Three-electrode stimulation is sufficient to steer the locus of activation in the plane of the tissue without recruiting additional unwanted neurons.

The biophysical priors used in fitting the linear-OR model and the direct estimation of noise in GPR make both models robust to experimental error in measurements of evoked activation probabilities. The fitted linear-OR model is able to overcome noisy data for two reasons. First, it implicitly assumes that activation probability increases monotonically and smoothly in every direction from the origin, removing some of the impact of individual noisy activation probabilities. Second, during model fitting, the parameter optimization is constrained to enforce that the activation probability corresponding to zero applied current on all electrodes is low (see Methods). In practice, these two considerations make the linear-OR model robust to the spike sorting errors, although a drawback of the model is that it does not explicitly estimate the noise in the data and is vulnerable to overfitting to noisy points in subsampled data. In contrast, one of the strengths of GPR is that it does explicitly estimate the noise in the data. During GPR hyperparameter optimization, the marginal log-likelihood of the data is maximized as a function of the noise variance, assuming that the observed data contain independent additive Gaussian noise. This makes GPR more robust to noisy data, although for aggressively subsampled data, the noise variance is more difficult to estimate.

Current steering with electrode triplets led to substantial increases in the ability to activate a single neuron selectively. This precision is important for future neuroengineering applications: although previous work suggested the possibility that single-neuron stimulation may be routinely achievable using individual electrodes in cell culture [34], stimulation selectivity is far more challenging in real neural circuits because cells are embedded in a dense neuropil instead of growing individually on the electrodes. Two methods were examined here for efficiently exploiting current steering for selective RGC activation. First, the linear-OR model was used to estimate selectivity landscapes by combining model fits to data from individual neurons to guide electrical stimulation choices. Though the model was accurate for the individual neurons, combining the model fits for several neurons was prone to propagation of fitting errors and generally captured only large-scale features of selectivity surfaces. Second, the more flexible GPR approach accurately captured details of the selectivity landscapes, particularly in regions of high selectivity. The uncertainty metric obtained from GPR also lent itself to the design of efficient calibration protocols to find selective stimuli. The tunability and robustness of GPR may allow for extensions to larger numbers of stimulating electrodes and to other neural interfaces.

Enhancing single-neuron selectivity of electrical stimulation could have a substantial impact on the ability to restore neural function compared to earlier approaches. The most widely used epiretinal implant for vision restoration [5] used large (200 µm) widely spaced (575 µm) electrodes, and produced coarse, irregular visual sensations that were not very useful to users. This is most likely because each electrode evoked activity indiscriminately in hundreds of RGCs of different types: in the central retina, RGC spacing is 5-30 µm [35]. In comparison, the present work used small (5-10 µm) closely spaced (30 µm) electrodes. At this scale, single-neuron stimulation is frequently but not always achievable [12–14], and selectivity can be enhanced using the current steering framework described here. With cellular-resolution control over a population of neurons, the neural code could be more accurately reproduced, potentially producing more faithful vision restoration.

Tools such as the linear-OR model and GPR+LSE, which allow for efficient calibration of selective electrical stimulation parameters, would be important for chronic *in vivo* and clinical settings in which implant movement and other factors can lead to significant changes in the electrical interface over time. The manifestation of such interface changes is an altered stimulation response surface for each neuron stimulated by a group of electrodes. The linear-OR model and GPR+LSE procedure presented here would permit rapid, repeated recalibration of the interface, finding new selective stimulation patterns to account for changes in the neural interface.

Due to the lack of precise knowledge of the positions of neurons relative to the MEA combined with the variety of linear and nonlinear responses to multi-electrode stimuli, empirical calibration would likely be required for a future implant to exploit cellular-resolution current steering, and the methods described here could reduce calibration times to clinically feasible timescales. For example, assuming a 10 ms interval between successive stimulations to avoid the neural refractory period, the GPR+LSE method required ∼1 minute of calibration time per electrode triplet. In selectivity landscapes with no selective regions, the algorithm terminated earlier, preventing unnecessary searches that would yield no selective stimulation patterns. Across the entire MEA used in this work with ∼1,000 distinct electrode triplets, assuming a maximum of 1 minute of calibration per triplet using GPR+LSE, the total calibration time is ∼16.5 hours, a timescale which could be feasibly divided over several clinical sessions to improve the precision of neural stimulation. The GPR+LSE framework exhibited a >1.5⨉ speedup compared to GPR+random sampling, potentially saving valuable clinic time.

Multi-electrode current steering may be useful in neuroengineering applications outside of the retina, and the calibration scheme developed here could provide a general framework for selective electrical stimulation of individual neurons. Specifically, the linear-OR model may be applicable to extracellular stimulation at single-neuron resolution in the brain, because the gray matter is predominantly composed of cell bodies and unmyelinated axons, as is the RGC layer of the retina. The efficient and generalizable methods presented here are particularly relevant for the larger-scale neural probes currently being developed [36]. As neural implants are increasingly used to restore and augment complex functionality, efficiently predicting and controlling the responses of individual neurons with patterned multi-electrode stimuli could become a cornerstone of brain-machine interfaces.

### Limitations of the study

The present work has several limitations and caveats:

● The proposed GPR+LSE algorithm was tested here by offline subsampling of exhaustively collected data rather than online in a true closed-loop experiment. In a true closed-loop approach, performing GPR and deciding which measurements to take next with LSE every iteration requires accurate, real-time spike sorting with limited data, motivating the need for one-shot spike detection algorithms for electrical stimulation in the presence of stimulation artifacts that do not require repeated trials of the same stimulus (A. Lotlikar, personal communication). Online spike sorting and GPR fitting will also add computation time in an experiment, but these temporal constraints can be alleviated by batching measurements between distinct electrode triplets across a large-scale MEA, i.e. performing computation for measurements obtained using one electrode triplet while simultaneously stimulating on another electrode triplet.
● The present study focused on parasol and midget RGCs. Though these cell types are numerically dominant in the primate retina and thus likely mediate much of high-resolution vision, there are many other cell types which are likely essential to natural vision, and it is possible that some of the findings would differ for those cell types.
● The multi-electrode stimuli used were presented simultaneously, ignoring the possible benefits of temporally modulated stimuli for achieving high stimulation selectivity.

## Methods

### Experimental setup

A custom 512-electrode system [9] was used to stimulate and record from RGCs in isolated rhesus macaque (*Macaca mulatta*) and human retinas. Macaque eyes were obtained from terminally anesthetized animals euthanized during the course of research performed by other laboratories. Human eyes were obtained from brain-dead organ donors through Donor Network West. All procedures were performed in accordance with institutional and national guidelines and regulations. Briefly, the eyes were hemisected in room light following enucleation, the vitreous was removed and the posterior portion of the eye was kept in darkness in warm (33–35°C), oxygenated, bicarbonate buffered Ames solution (Sigma). Then, patches of retina ∼2 mm on a side were isolated under infrared light, placed RGC side down on the microelectrode array (MEA), and superfused with oxygenated Ames solution. Central retinal preparations corresponded to regions with 1.5–4.5 mm (4.5–20°) eccentricity, while peripheral preparations ranged from 5–12 mm in eccentricity (22–56° temporal equivalent). The MEA was composed of 512 electrodes (diameter 5-10 µm) in an isosceles triangular lattice with center-to-center distance between adjacent electrodes either 30 µm or 33.5 µm, covering an area of 0.43 mm^2^.

Within an experiment, platinization of the electrodes produced relatively uniform noise: the RMS noise had a ∼6% standard deviation across electrodes. The platinized electrodes had an impedance of ∼200 kΩ at 1 kHz, a typical frequency for recording action potentials, leading to a negligible voltage divider effect between the electrodes and the input impedance of each electrode’s amplifier (∼100 MΩ), resulting in < 1% signal attenuation. A platinum wire encircling the recording chamber (∼1 cm diameter) served as the distant return electrode. Voltage recordings were band-pass filtered between 43 and 5,000 Hz and sampled at 20 kHz. Spikes from individual RGCs in the voltage recordings obtained with visual stimulation of the retina were identified and sorted using Kilosort [37].

### Visual stimulation and cell type classification

To identify the retinal ganglion cell types recorded, the retina was visually stimulated with a dynamic white noise stimulus, and the spike-triggered average (STA) stimulus was computed for each RGC, as previously described [38,39]. The STA summarizes the spatial, temporal, and chromatic properties of light response. In particular, clustering on the spatial (receptive field size) and temporal (time course) components of the STA was performed to identify distinct cell types, as previously described [40]. This process permitted rapid identification of ON and OFF parasol and midget RGCs, the four numerically dominant RGC types in primates.

### Electrical images

The electrical image (EI) represents the average spatiotemporal pattern of voltage deflections produced on each electrode of the array during a spike from a given cell [41]. Electrical images were calculated from data recorded during visual stimulation and served as spatiotemporal templates for the spike waveforms of the cells to be detected during electrical stimulation.

The spatial positions of relevant cell compartments (axon, soma, dendrites) of a recorded cell relative to the electrode array were estimated using the electrical image. The shape of the spike waveform from the EI on an electrode was used for compartment identification: a triphasic waveform was taken to indicate recording from the axon, a biphasic waveform with a positive first phase was taken to indicate recording from the dendrites, and a biphasic waveform with a negative first phase was taken to indicate recording from the soma [41].

### Electrical stimulation

Electrical stimulation was provided through one, two or three electrodes simultaneously while recording RGC activity from all 512 electrodes. Constant-current stimulation was used to correct for nonuniformities in the electrode impedance due to the electroplating procedure, ensuring that the injected current matched the desired current levels. Stimulating with a given electrode consisted of passing a charge-balanced, triphasic current pulse through that electrode. The cathodic-peak stimulus consisted of a triphasic pulse with anodal/cathodal/anodal phases with relative current amplitudes 2:-3:1 and duration of 50 μs per phase (150 μs total). The anodic-peak stimulus was similar but with the polarity of the phases reversed to be cathodal/anodal/cathodal. All stimuli were delivered in 20 repeated trials at combinations of currents from 20 linearly spaced second phase amplitudes between −1.8 μA and 1.8 μA. As a result, for three-electrode stimulation, 8000 unique current combinations were tested. The ordering of the stimulation patterns was chosen pseudo-randomly, restricted so that each stimulating electrode or group of electrodes was far from the previous stimulating electrodes to avoid stimulating the same cell(s) in rapid succession. A time delay of 10 ms was enforced between successive stimulations to ensure that the neural relative refractory period had elapsed and that successive trials could be considered independent. The stimulating electrodes were chosen based on the highest SNR of recorded spikes from the target cell as well as the geometric positioning of the electrodes relative to the target cell.

### Responses to electrical stimulation

The spikes recorded during electrical stimulation were analyzed using a custom template matching approach [12]. First, the electrical image (EI) of each cell was calculated from visual stimulation data to serve as a template for the spike waveform of each cell to be detected during electrical stimulation. In the electrical stimulation data, an automated algorithm separated spikes from the electrical artifact by clustering traces obtained from repeated trials of the same electrical stimulus and comparing the waveform difference between these clusters to the EI templates obtained from visual stimulation.

The neurons considered during electrically-evoked spike sorting for a group of stimulating electrodes were determined by the SNR of recorded spikes on the same electrodes. Specifically, for a group of stimulating electrodes, if any of the electrodes recorded a spiking amplitude greater than or equal to 2σ for a given cell, with σ the standard deviation of noise on the electrode, that cell was considered during electrically-evoked spike sorting. Larger recorded spike amplitudes are known to be correlated with greater stimulation sensitivity using the same electrodes, as detailed in [42], and this threshold of 2σ was chosen conservatively to ensure that all relevant neurons were considered for each group of stimulating electrodes. For lower SNR levels, electrically-evoked spike sorting is not reliable, and the neurons were often not activated by stimulation [42].

Response probabilities for each cell were calculated for all current combinations passed through the three electrodes. For plotting experimental stimulation responses, only combinations causing response probability 0<p<1 were used to more clearly display the shape of the response in the three-dimensional space of currents.

### Biophysical model

The biophysical model of a spiking parasol RGC was developed in the NEURON [24] simulation environment. Full details are provided in [25]. Briefly, the biophysical properties of the RGC cable model used were adapted from [30] with modifications to the temperature of the simulation environment and the membrane dynamics as detailed in [43]. Additionally, the narrow axon compartment was removed, and the remaining compartment diameters were increased, which allowed for more stable model behavior when simulating electrical stimulation with multiple electrodes. All code used to build the RGC model is online and available at: https://github.com/ramanvilkhu/rgc_simulation_multielectrode

### Linear-OR model fitting

The linear-OR model is described by the predicted probability given in Equation 1. The model weights were fitted using maximum likelihood estimation (MLE) considering the cell spiking responses for each current vector as binary outputs *Y_j_* drawn i.i.d. from a Bernoulli distribution with parameter *p* = *p*(*x_j_*). The negative log-likelihood for this Bernoulli distribution with N observations (every independent trial at every current level tested) is given in Equation 3:

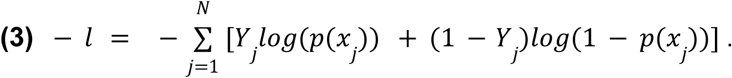

Equation 3 can be equivalently written in terms of a weighted sum with the weights being the number of repeated trials at every current level, reducing computation time.

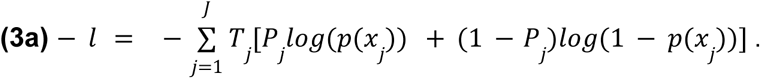

Here, *P_j_* is the empirically measured activation probability at current level *x_j_* and *T_j_* is the number of repeated trials performed at *x_j_* to calculate *P_j_*. This negative log-likelihood can be shown to be non-convex for the form of *p*(*x*) given in Equation 1. In practice, solutions within parameter constraint bounds were found using second-order methods and regularization. The L-BFGS-B optimizer [44] was used for constrained minimization of Equation 3a with constraints on the bias parameters *w*_i,0_ to enforce that the model fit produced a sufficiently low probability (0.01) with *x* = 0 and constraints such that the weight parameters *w*_i,k_ ≤ 100. An L2 regularization penalty (λ = 0.5) was also added during optimization.

The process of fitting the linear-OR model also involved choosing a hyperparameter: the number of sites (*m* in Equation 1), which in turn controlled the allowed degree of nonlinearity of fitted surfaces. The value *m* = 1 corresponded to logistic regression. In practice, a linear model with *m* = 1 reduces to a null model with only an intercept and no weightings on the currents when fitted to double-planar and nonlinear data. A fixed value of *m* = 8 was used to account for significant nonlinearities in the three-electrode data. For fitting to data from biophysical simulations of four-electrode stimulation, *m* = 10 was used. The goodness of fit was quantified using the McFadden pseudo-R^2^ metric [45], one of several measures of goodness of fit commonly used for binary regression tasks. The definition of the McFadden pseudo-R^2^ is given in Equation 4:

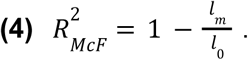

Here, *l*_m_ is the log-likelihood of the fitted model with number of sites *m* and *l*_0_ is the log-likelihood of the null model which features only an intercept and no weighting on the input currents *x*. All code used for fitting the linear-OR model can be found at: https://github.com/pvasired/Vasireddy2025

### Gaussian Process Regression and Level Set Estimation

Gaussian process models are nonparametric models describing a probability distribution over possible functions (see [26]). Gaussian process regression (GPR) is an established method for using Gaussian process models to fit a set of observations and predict outputs on novel inputs (see [27]).

In this application, the inputs are the multi-electrode stimulation amplitude vectors *x*. Assuming there are *K* cells *c*_1_,…,*c*_K_ stimulated by stimulus *x* at different activation probabilities *p*_1_,…,*p*_K_, an aggregate selectivity index can be constructed as in Equation 5:

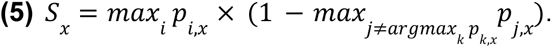

This aggregate metric yields the highest selectivity achievable for any cell using stimulus *x*, agnostic to which cell is selectively stimulated. This index ranges from 0 to 1, with 1 representing perfect selectivity. Rather than performing GPR directly on the selectivity values from Equation 5 that are bounded between 0 and 1, which would introduce additional complexity to modeling, a logit transformation of the aggregate selectivity index was performed to produce an unbounded real number, 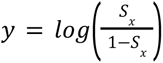. GPR was then performed on the collection of pairs {(*x*, *y*)} across stimulus amplitude vectors. To predict activation probabilities of individual neurons (as in Fig. 4D-F, Fig. 5D-F), the logit-transformed activation probabilities for each cell were used, 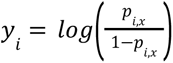.

Standard (i.e., non-sparse) GPR was implemented in the GPyTorch framework [46]. A constant mean function prior was assumed, and the RBF kernel was used for the covariance. Hyperparameter optimization of the RBF kernel length scale and the noise variance was performed using the AdamW optimizer [47]. The GPR predictions remained in logit (log-odds) units, but they could be transformed back to the bounded selectivity index using a sigmoid transformation.

With the fitted model from GPR on subsampled data, predictions were made across the entire input space of both the posterior mean function *μ* and variance *σ*. Using these predictions, the space of possible inputs was efficiently explored to accurately estimate the high selectivity regions. A variant of the GP-UCB algorithm [48] with similarities to methods from level set estimation (LSE; see [49]) was implemented to encourage exploration of regions that could yield high selectivity while preventing oversampling of regions known with high certainty to yield high selectivity. In the conventional GP-UCB algorithm, the next measurements to take are decided by choosing inputs that maximize the *upper confidence bound UCB = μ* + *βσ* of the current GPR fit, with *β* a hyperparameter controlling how wide of a confidence bound is considered. Larger values of *β* in GP-UCB will encourage exploration of unknown regions by strongly weighting uncertainty in the GPR predictions, while smaller values of *β* will encourage concentrated sampling of regions known to yield high posterior mean function values (see [48] for a fuller discussion of choosing *β*). In this implementation, the *lower confidence bound LCB = μ* - *βσ* is also used to exclude potential regions to sample. Specifically, during the fitting process, if *LCB > L* for a given unmeasured input, that input is removed from the set of possible inputs to sample next. Of the remaining set of allowed inputs to sample, the next batch of measurements is chosen randomly amongst inputs which have *UCB > L*. In this implementation, *β* = 2, *L* = 3, and a batch size of 1000 measurements per iteration were used. Intuitively, this algorithm prevents sampling points which are nearly certain to yield high selectivity (*LCB > L*) and do not need further exploration while also sampling unexplored regions which could yield high selectivity (*UCB > L*). All code used here for GPR and LSE can be found at: https://github.com/pvasired/Vasireddy2025

## Acknowledgements

National Science Foundation Grant No. 1828993 (PKV), ALS Association Milton Safenowitz fellowship (NPS), Excellence Initiative – Research University for the AGH University (PH), Research to Prevent Blindness Stein Innovation Award, Wu Tsai Neurosciences Institute Big Ideas, NIH NEI R01-EY021271, and NIH NEI P30-EY019005 (EJC). We thank Sasi Madugula, Vincent Zhuang, Scott Linderman, and the Stanford Artificial Retina team for experimental contributions and useful discussions.

## Author Contributions

P.K.V. and E.J.C. conceived of the multi-electrode stimulation experiments and the approach for studying single-neuron responses to current steering. P.K.V., R.S.V., and A.R.G. designed the multi-electrode stimulation experiments and collected the experimental data. P.K.V., A.L., and E.J.K. performed the downstream analysis and empirical modeling. P.K.V. and J.B.B. conceived of the specific mathematical framing of the linear-OR model. A.L. and J.B.B. designed algorithms for spike sorting. R.S.V. implemented the biophysical simulations. A.J.P., M.R.H., C.B., and Av.S. assisted with experimental data collection. P.H., Al.S., and A.M.L. designed and constructed the microelectrode array system for stimulation and recording. P.K.V. prepared figures with input from R.S.V., A.J.P., P.H., E.J.K., N.P.S., and E.J.C. P.K.V., N.P.S., and E.J.C. wrote the manuscript with inputs from all authors.

---

**Extended Data Figure 1.**
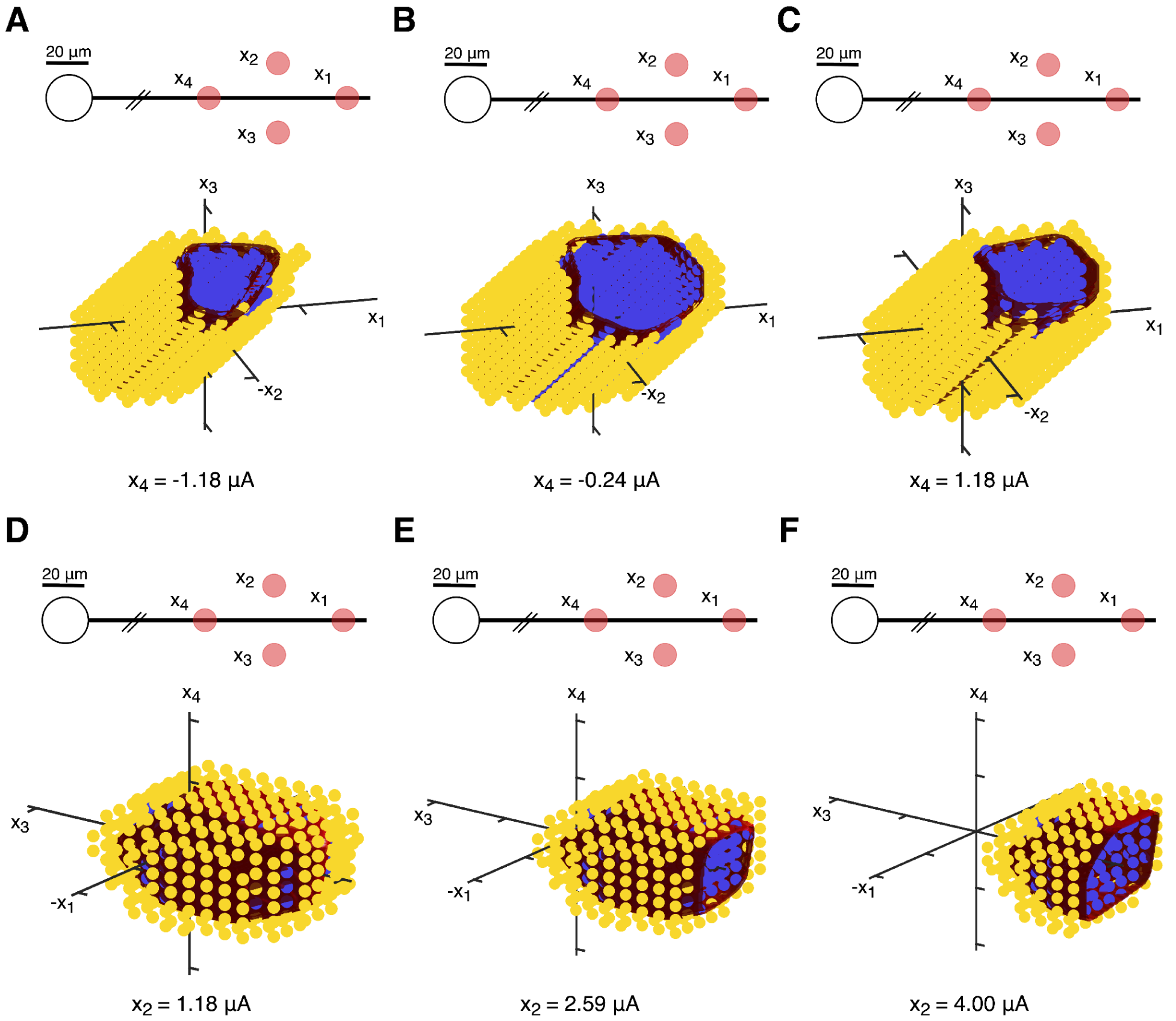
Extension of linear-OR model fitting to four-electrode stimulation responses in biophysical simulations. Nonlinear four-electrode stimulation biophysical simulation data with linear-OR model 50% surface for *m* = 10 activation sites. The linear-OR model fit had a McFadden pseudo-R^2^ of 0.86, indicating an excellent fit (see Methods). Slices with freely varying amplitudes on three electrodes and fixed fourth electrode amplitude are shown. ***A-C,*** Stimulation responses along three-electrode slices with fixed amplitude fourth electrode along the axon, for various fixed amplitude levels on the fourth electrode. ***D-F,*** Stimulation responses along three-electrode slices with fixed amplitude fourth electrode transversely offset from the axon, for various fixed amplitude levels on the fourth electrode.

**Extended Data Figure 2.**
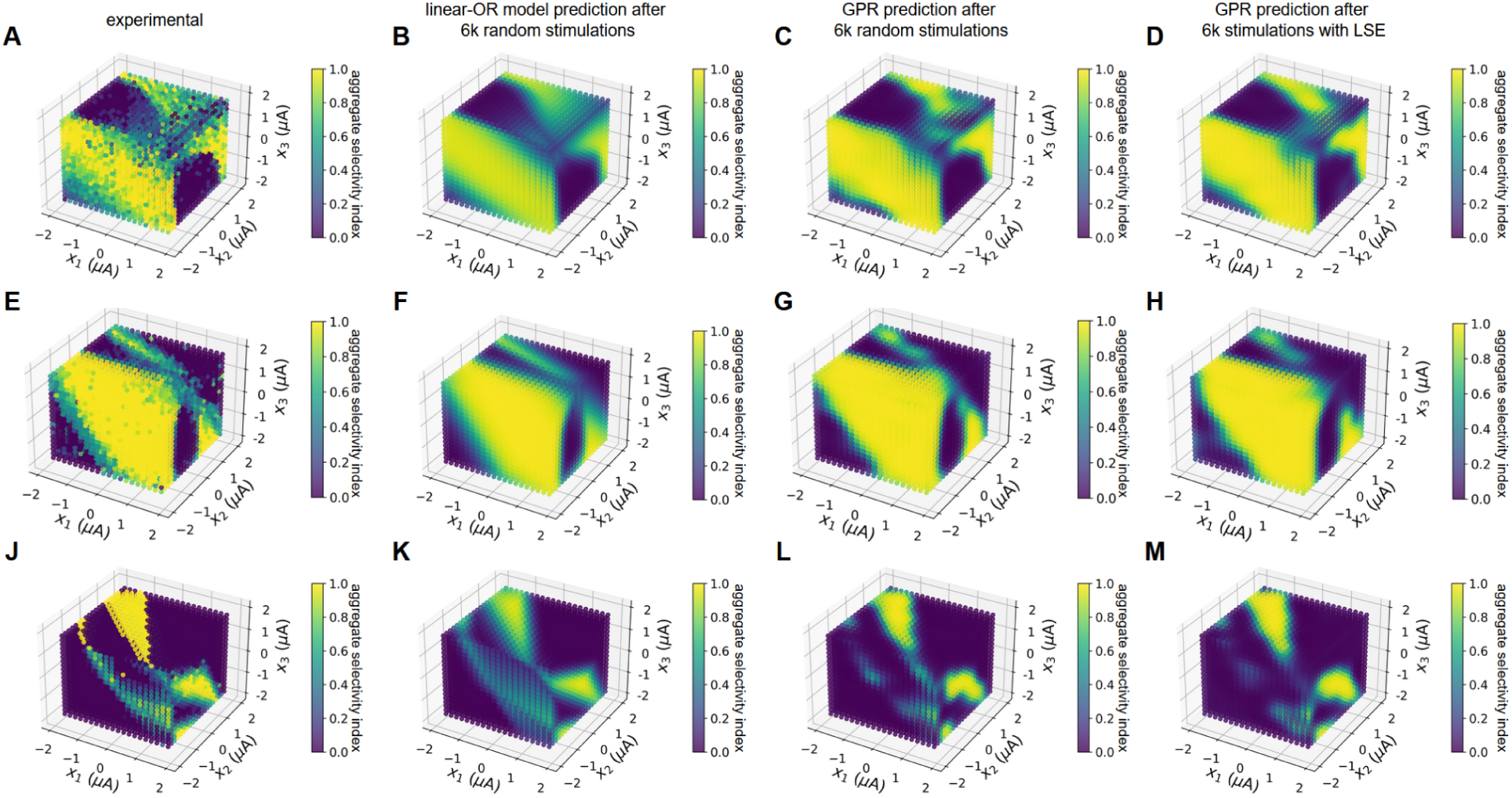
Gaussian process regression (GPR) and linear-OR model for estimation of aggregate selectivity profiles. First column shows experimentally measured aggregate selectivity profiles from full data (160,000 stimulations), second column shows linear-OR model prediction after fitting to subsampled data from each cell and computing the aggregate selectivity index, third column shows GPR mean prediction after performing 6,000 random calibrating stimulations, and fourth column shows GPR mean prediction after performing 6,000 calibrating stimulations guided by the GPR+LSE sampling procedure. Each row represents a distinct electrode triplet from a particular retinal preparation. ***A-D,*** Macaque peripheral RGCs (2 cells aggregated). ***E-H,*** Macaque central RGCs (5 cells aggregated). ***J-M,*** Macaque peripheral RGCs (4 cells aggregated).

